# Adult Canine Intestinal Derived Organoids: A Novel *In Vitro* System for Translational Research in Comparative Gastroenterology

**DOI:** 10.1101/466409

**Authors:** Lawrance Chandra, Dana C Borcherding, Dawn Kingsbury, Todd Atherly, Yoko M Ambrosini, Agnes Bourgois-Mochel, Wang Yuan, Michael Kimber, Yijun Qi, Qun Wang, Michael Wannemuehler, N Matthew Ellinwood, Elizabeth Snella, Martin Martin, Melissa Skala, David Meyerholz, Mary Estes, Martin E. Fernandez-Zapico, Albert E. Jergens, Jonathan P Mochel, Karin Allenspach

## Abstract

**Background:** Large animal models, such as the dog, are increasingly being used over rodent models for studying naturally occurring diseases including gastrointestinal (GI) disorders. Dogs share similar environmental, genomic, anatomical, and intestinal physiologic features with humans. To bridge the gap between currently used animal models (e.g. mouse) and humans, and expand the translational potential of the dog model, we developed a three dimensional (3D) canine GI organoid (enteroid and colonoid) system. Organoids have recently gained interest in translational research as this model system better recapitulates the physiological and molecular features of the tissue environment in comparison with two-dimensional cultures.

**Results:** Organoids were propagated from isolation of adult intestinal stem cells (ISC) from whole jejunal tissue as well as endoscopically obtained duodenal, ileal and colonic biopsy samples of healthy dogs and GI cases, including inflammatory bowel disease (IBD) and intestinal carcinomas. Intestinal organoids were comprehensively characterized using histology, immunohistochemistry, RNA *in situ* hybridization and transmission electron microscopy, and organoids mimicked the *in vivo* tissue environment. Physiological relevance of the enteroid system was defined using functional assays such as Optical Metabolic Imaging (OMI), the Cystic Fibrosis Transmembrane Conductance Regulator (CFTR) function assay, and Exosome-Like Vesicles (EV) uptake assay, as a basis for wider applications of this technology in basic, preclinical and translational GI research.

**Conclusions:** In summary, our findings establish the canine GI organoid systems as a novel model to study naturally occurring intestinal diseases in dogs and humans. Furthermore, canine organoid systems will help to elucidate host-pathogen interactions contributing to GI disease pathogenesis.

## Background

Rodent models, especially the mouse, have been extensively used to study gastrointestinal (GI) diseases due to cost effectiveness, ethical considerations and the easy accessibility to genetically engineered technology. Despite the wide use of mouse models in biomedical research, the translational value of mouse studies for human disease remains controversial [1]. In addition, mice and other rodents often fail to adequately represent the human condition, as well as drug response in toxicity and efficacy studies [2, 3]. Given the high failure rate of drugs from discovery and development through the clinical trial phase (i.e. more than 90%), there is now a critical need for better animal models for preclinical studies [4].

Large animal models, such as the dog, are typically more representative than mice as they have a relatively large body size, longer life span, more closely resemble human GI physiology and develop spontaneous, analogous diseases including inflammatory bowel disease (IBD) and colorectal cancer (CRC) [4]. Dogs have been used as an animal model for human health and disease from the ancient to the modern era [5, 6]. The dog is still considered to be a superior non-rodent mammalian animal model for pharmaceutical research, and is preferred by the FDA for initial safety data of drugs for human use [7]. Although the dog has contributed immensely to the advancement of medical knowledge in the past, the use of the dog in medical research has declined in recent years due to the emotional perceptions among the public and ensuing ethical issues with canine research [5]. The canine GI organoids arose as a model to bridge the gap in the drug development pipeline by providing a more representative *in vitro* model to test drug efficacy and toxicity in preclinical studies, as well as an innovative screening tool in drug discovery, while also reducing the number of animals needed for *in vivo* studies [2, 4, 8]. Thus, the ultimate goal of our research is to culture canine intestinal organoids from diseased dogs to develop better therapeutic strategies and personalized medicine for both animal and human health.

Stem cell-derived 3D organoids have emerged as a cutting edge cell culture technology to study the developmental biology of the intestines, brain, stomach and liver [9–12], drug discovery and toxicity screening [4], drug testing for personalized medicine [4,13], infectious disease biology of viruses [14, 15], and regenerative medicine [16]. Organoids are collections of organ-specific cell aggregates derived from either primary tissue or stem cells that are capable of organ-like functionality in an *in vitro* environment [17, 18]. The 3D organoid model better reproduces the *in vivo* biology, structure, and function, as well as genetic and epigenetic signatures of original tissues, unlike widely used two dimensional (2D) cell monolayer models that utilize cancer and immortalized cell-lines [4, 19–21].

Organoids are developed from either embryonic or induced pluripotent mesenchymal-derived stem cells (iPSC) or organ-specific adult stem cells (ASC) [17, 19]. Organoids derived from ASCs are generated without genetic transduction by transcription factors, unlike organoids derived from iPSCs [19], thus providing a more physiologically relevant *in vitro* model than iPSC-derived organoids. ASC-derived organoids are a functional model that can be differentiated to replicate the *in vivo* adult environment, and can be safely transplanted into animals and humans [22–23]. In addition, adult intestinal stem cell (ISC)-derived organoids have recently gained attention as a model to understand how the intestinal epithelia interact with the gut microbiome to modulate GI health and disease, for the study of infectious diseases of the GI tract, and as a drug screening tool for personalized medicine in diseases such as cystic fibrosis (CF) [13, 24].

In this study, we have developed 3D canine intestinal organoids from healthy dogs and dogs with GI diseases, including IBD and intestinal carcinomas. Intestinal organoids, propagated from leucine-rich repeat containing G protein-coupled receptor 5 (Lgr5)-positive stem cells located in intestinal crypts, are termed ‘enteroids’ or ‘colonoids’, depending on the anatomic region of origin (i.e. small vs. large intestine). Jejunal 3D enteroids were characterized from healthy dogs using histopathology, immunohistochemistry (IHC), RNA *in situ* hybridization (RNA-ISH) and transmission electron microscopy (TEM) and compared to full-thickness jejunal tissues to prove the reproducibility and translatability of this *in vitro* model. The physiological relevance of the canine enteroid system was further demonstrated using functional assays, including Optical Metabolic Imaging (OMI), the Cystic Fibrosis Transmembrane Conductance Regulator (CFTR) function assay, and the Exosome-Like Vesicles (ELVs) uptake assay. In summary, 3D canine enteroids are a relevant *in vitro* animal model with wide applications in veterinary and translational biomedical research: (1) to perform mechanistic studies for basic GI research, (2) for applied preclinical drug permeability, efficacy, and safety testing; (3) for personalized medicine in animal health and (4) for preclinical research prior to *in vivo* clinical trials in human patients.

## Results

### Development of 3D cultures of canine enteroids and colonoids

Using isolated canine adult intestinal stem cells (ISC), we developed 3D intestinal enteroids and colonoids from 28 healthy and 20 diseased dogs, including dogs diagnosed with IBD (N=8) and intestinal mast cell tumor (N=2). Enteroids and colonoids were collected from different intestinal segments, including the duodenum, jejunum, ileum and colon, and were maintained for up to over 20 passages. A summary of the demographics of dogs used for canine intestinal stem cell isolation, culture and maintenance is presented in Table 1. Intestinal stem cell isolation and enteroid and colonoid maintenance followed a modified version of the procedure previously described by Saxena *et al* for human organoids, and the defined media included Wnt3a, R-spondin-1, and Noggin, as well as inhibitors of Rock and GSK3β for the first 2-3 days of culture [25]. Figure 1 shows the phase contrast images of fully differentiated 6-7 day-old duodenal, jejunal, and ileal enteroids as well as colonoids from healthy dogs and dogs with IBD. There were no major morphological differences in the structures of enteroids and colonoids from healthy dogs and dogs with IBD, as visualized by bright field microscopy. We created a bio-archive of all enteroids and colonoids for future use in basic and applied research. This resource, which is to date the largest collection of 3D enteroids and colonoids from healthy and diseased dogs, will be made available to the wider scientific community (Table 1).

**Figure 1.**
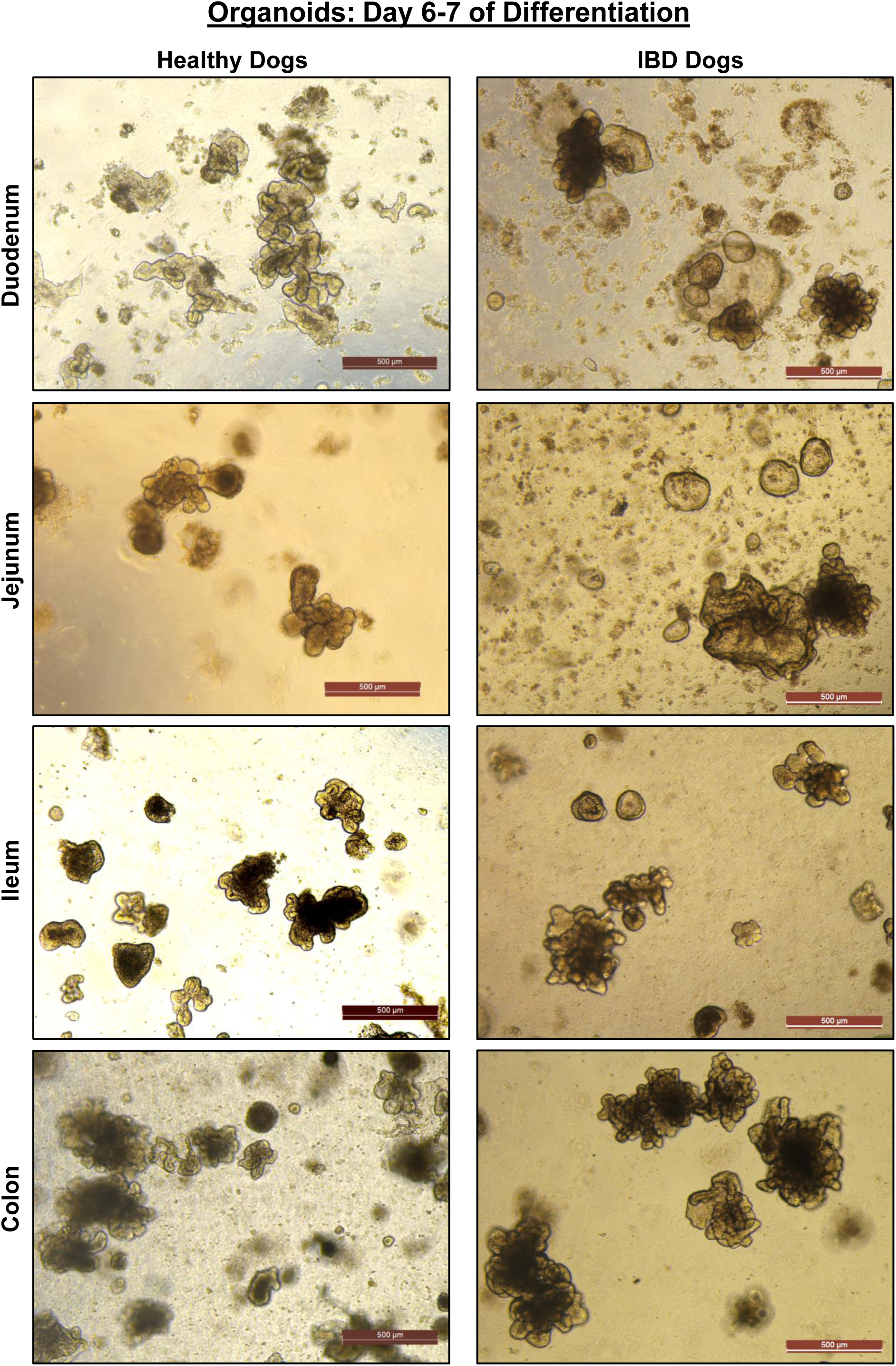
Similar structure and differentiation of canine intestinal 3D organoids, both enteroids and colonoids, derived from different intestinal compartments of healthy dogs and dogs with Inflammatory Bowel Disease (IBD). Representative phase contrast images of fully differentiated 6-7 day-old duodenal, jejunal, and ileal enteroids as well as colonoids from healthy dogs and dogs with IBD. There were no major difference in the structures of enteroids and colonoids from healthy dogs and dogs with IBD.

**Table 1.** Details of dogs used for isolation, propagation and preservation of canine 3D intestinal organoids.

|  |  |  |  |  | # of vials archived |  |  |  |
| --- | --- | --- | --- | --- | --- | --- | --- | --- |
| S. No | Type of Dog | # of Dogs | Tissue type | Tissue site/location | Duodenum | Jejunum | Ileum | Colon |
| 1 | Healthy | 20 | Whole tissue | 15 dogs: Jejunum |  | 78 |  |  |
|  |  |  |  | 5 dogs: Duodenum, Ileum, Colon | 2 |  | 14 | 15 |
| 2 | Nutrition Dogs: |  |  |  |  |  |  |  |
| 2.a | Pre treatment | 8 | Biopsy | Duodenum, Ileum, Colon | 23 |  | 30 | 26 |
| 2.b | Post treatment | 8 | Biopsy | Duodenum, Ileum, Colon | 39 |  | 56 | 58 |
| 3 | IBD Dogs | 9 | Biopsy | Duodenum, Ileum, Colon | 8 |  | 13 | 14 |
| 4 | Lysosyme storage mutation | 1 | Whole tissue | Duodenum, Ileum, Colon |  |  | 4 | 4 |
| 5 | Tumor | 2 | Biopsy |  | 5 |  | 2 | 2 |

### Morphologic characterization demonstrates high resemblance of canine organoids to original tissue

Histology of enteroids on Days 3, 6 and 9 of differentiation is shown in representative images of H&E tissue sections (Fig 2). Day 3 enteroids have undifferentiated cyst-like structures embedded in pink-colored matrigel 3D matrix, whereas Days 6 and 9 enteroids present both crypt and villi-like structures embedded in matrigel. Dark purple staining bodies in the enteroid lumen represent epithelial cells that have undergone apoptosis in Day 9 tissue sections. For comparison, the H&E tissue sections of full-thickness jejunal mucosa reveal normal tissue architecture.

**Figure 2.**
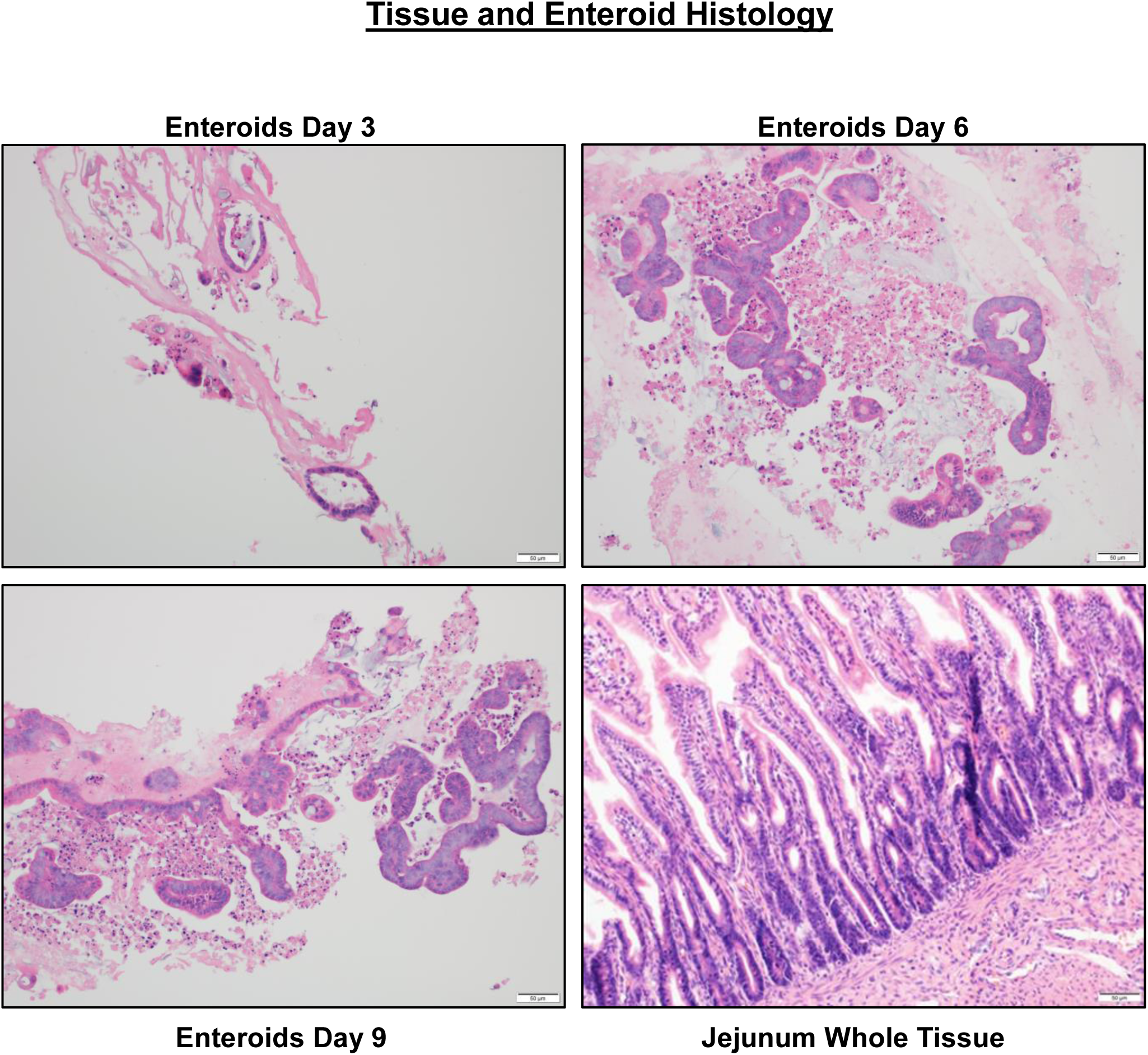
Characterization of canine jejunal tissue and jejunal enteroid histology shows similarities of epithelial structure. Histological images of hematoxylin and eosin (H & E) staining show the development and differentiation of canine enteroids at 3, 6 and 9 days after isolation or passaging. Spheroid-like epithelial structures are visualized in 3 day enteroids compared to crypt-villi epithelial structures on the 6th and 9th days. In the whole jejunal tissue there arecrypt-villi epithelial structures, non-epithelial cell types are also seen.

To identify changes in the ultrastructural and morphometric details of enteroids during differentiation on Days 3, 6, and 9, we used TEM, which allowed longitudinal assessment of morphologic features. TEM revealed that differentiation of cells also occurs on an ultrastructural level. In jejunum tissue and enteroids, we visualized morphologic features of differentiated epithelial cells, including electron-lucent cytoplasmic vacuoles in mucus-producing goblet cells, and electron-dense perinuclear (neurosecretory) granules in enteroendocrine cells (Fig 3a). In addition, TEM showed the presence of microvilli at the apical border of epithelial cells, which increased throughout enteroid differentiation (Fig 3a). Microvilli are cellular membrane protrusions of absorptive enterocytes containing different populations of brush border enzymes involved in absorption, secretion and cellular adhesion [26]. Evidence of epithelial differentiation also involves increasingly adherent inter-epithelial structures, including adherens junctions (AJ), tight junctions (TJ) and desmosomes, and these were identified in both canine enteroids and in native jejunum (Fig 3b). On day 3, the developing tight junction structure had a dilated paracellular space between epithelial cells, with the space diminishing on day 6 and no longer apparent by day 9 (Fig 3b).

**Figure 3.**
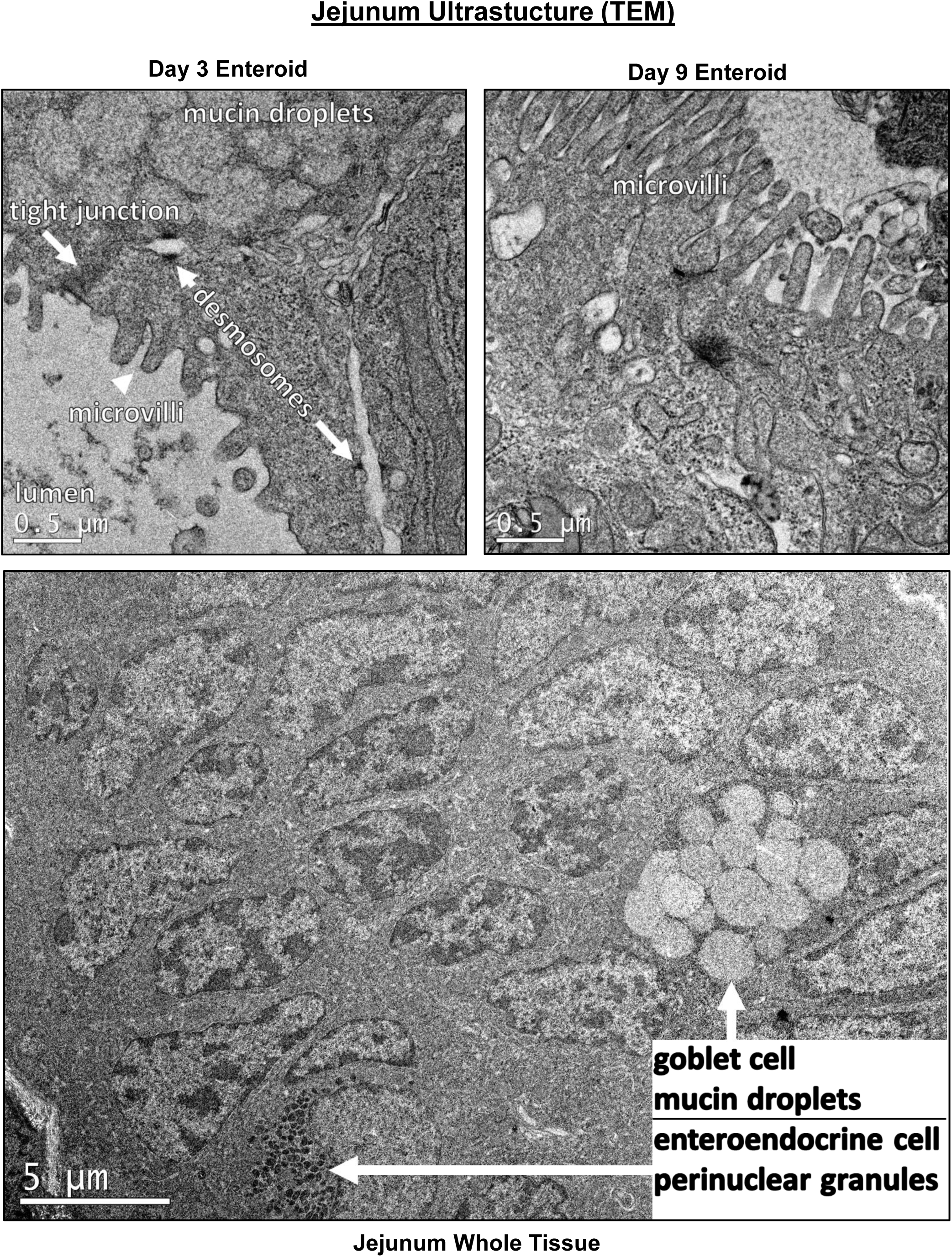
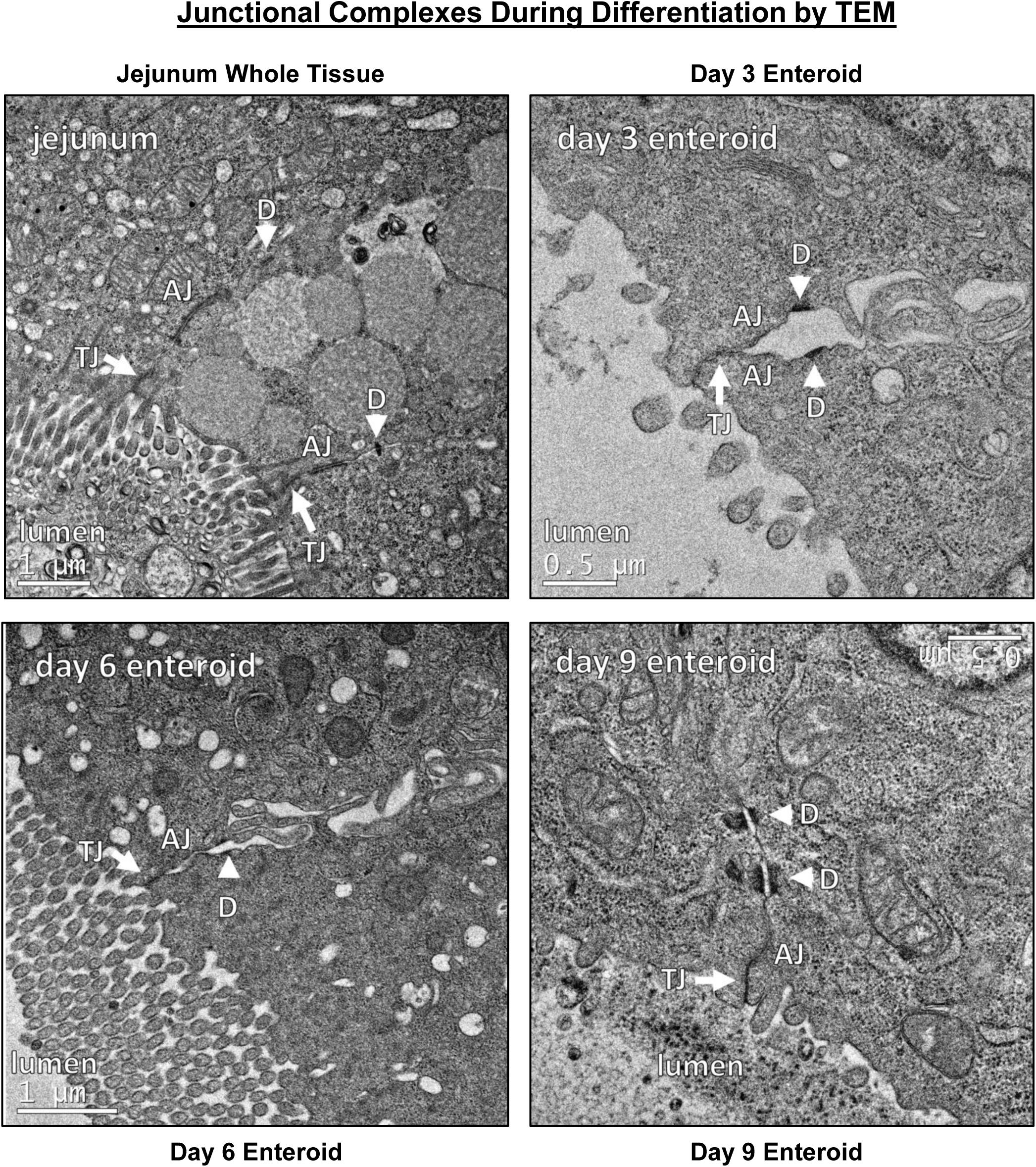
Ultrastructural features of differentiated enteroids mimic intact small intestinal tissue. **a** Representative transmission electron micrographs (TEM) of canine jejunum enteroids and whole tissue show features of cellular differentiation such as apical microvilli, electron-lucent cytoplasmic vacuoles (i.e. mucus), and electron-dense perinuclear granules (i.e. neurosecretory granules), consistent with development of absorptive enterocytes, goblet cells and enteroendocrine cells, respectively. **b** TEM of canine organoids show the progressive development (days 3-9 of differentiation) of intercellular structures important for intestinal barrier function. Adherens junction (AJ), tight junction (TJ) and desmosomes (D) structures are seen in both canine enteroids and
in native jejunum. On day 3, the developing tight junction had dilated paracellular space adjacent to tight junctions; however, the paracellular spaces were smaller on day 6, and no longer apparent by day 9.

### Molecular marker analysis of canine organoids

Although enteroids from several locations and colonoids were obtained from both healthy and diseased dogs, we first performed phenotypic characterization of jejunal enteroids due to their importance in nutrient absorption and drug permeability. For IHC, we used antibodies to identify cell surface expression of epithelial cells and their lineage proteins (Pan-Keratin for epithelial cells, Chromogranin A for enteroendocrine cells, and PAS for goblet cells), mesenchymal cells (Vimentin and Actin), Paneth cells (Lysozyme) and immune cells (c-Kit and CD3) in jejunal enteroids compared to full-thickness jejunal tissues. IHC markers for epithelial cells and their lineage, such as Keratin, Chromogranin A and PAS had positive staining in both intact jejunal tissue and jejunal enteroids (Fig 4 and Table 2). Keratin staining was absent in the lamina propria region of whole jejunal tissue, confirming the specificity of the epithelial marker in canine intestines [27]. Lysozyme, a marker of Paneth cells, was absent both in enteroids and jejunal epithelium (Table 2), which is consistent with previous reports showing that canine intestines lack Paneth cells [28]. Conversely and as expected, jejunal enteroids did not express markers for mesenchymal cells (Vimentin and Actin) or immune cells (c-Kit and CD3, a T cell marker) as they only contained epithelial cells (Table 2), whereas intact jejunal tissues robustly expressed all mesenchymal and immune cell markers in the lamina propria (Table 2).

**Table 2.** Characterization of canine jejunal tissue and jejunal enteroids using IHC.

| **Tissue/Enteroid** | **Stain /Marker** | **Result** | **Location** |
| --- | --- | --- | --- |
| Enteroid | Keratin / Epithelium | Pos | Epithelium |
| Jejunum | Keratin / Epithelium | Pos | Epithelium |
| Enteroid | PAS /Goblet | Pos | Goblet cells |
| Jejunum | PAS /Goblet | Pos | Goblet cells |
| Enteroid | Chromogranin A /<br>Enteroendocrne | Pos | Epithelium |
| Jejunum | Chromogranin A /<br>Enteroendocrne | Pos | Epithelium |
| Enteroid | Vimentin /Mesenchymal | Neg |  |
| Jejunum | Vimentin /Mesenchymal | Pos | Submucosa, lamina propria<br>vessels, muscularis |
| Enteroid | Actin /Mesenchymal | Neg |  |
| Jejunum | Actin /Mesenchymal | Pos | Muscularis, lamina propria<br>vessels |
| Enteroid | c-Kit / Leukocyte | Neg |  |
| Jejunum | c-Kit / Leukocyte | Pos | Resident leukocytes |
| Enteroid | T cell/ T lymphocyte | Neg |  |
| Jejunum | T-cell/ T lymphocyte | Pos | Intraepithelial and lamina<br>propria lymphocytes |
| Enteroid | Lysozyme / Paneth cell | Neg | Epithelium |
| Jejunum | Lysozyme / Paneth cell | Neg | Epithelium |

**Table 3.**
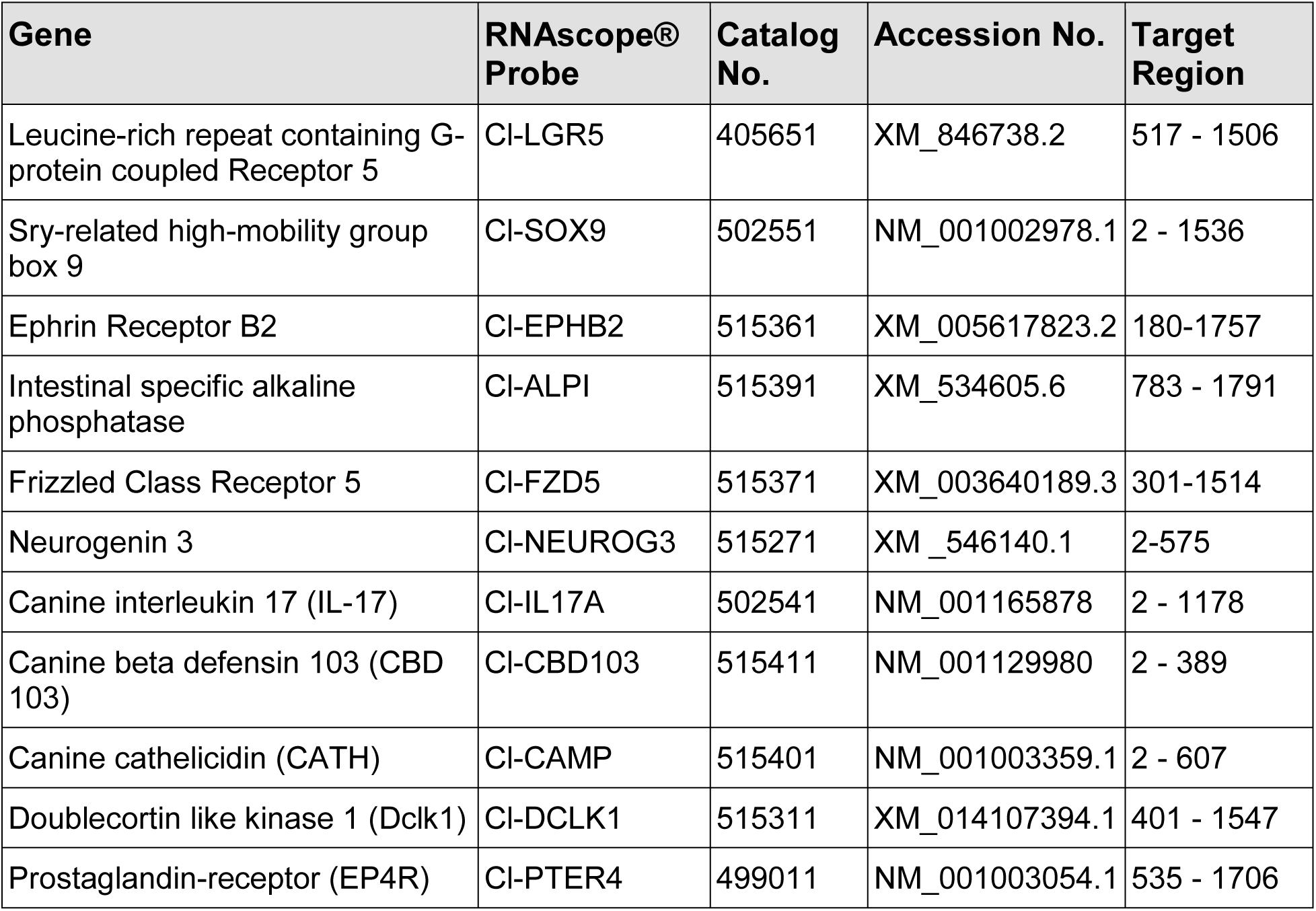
RNA *In Situ* Hybridization (ISH) Probe Details.

**Figure 4.**
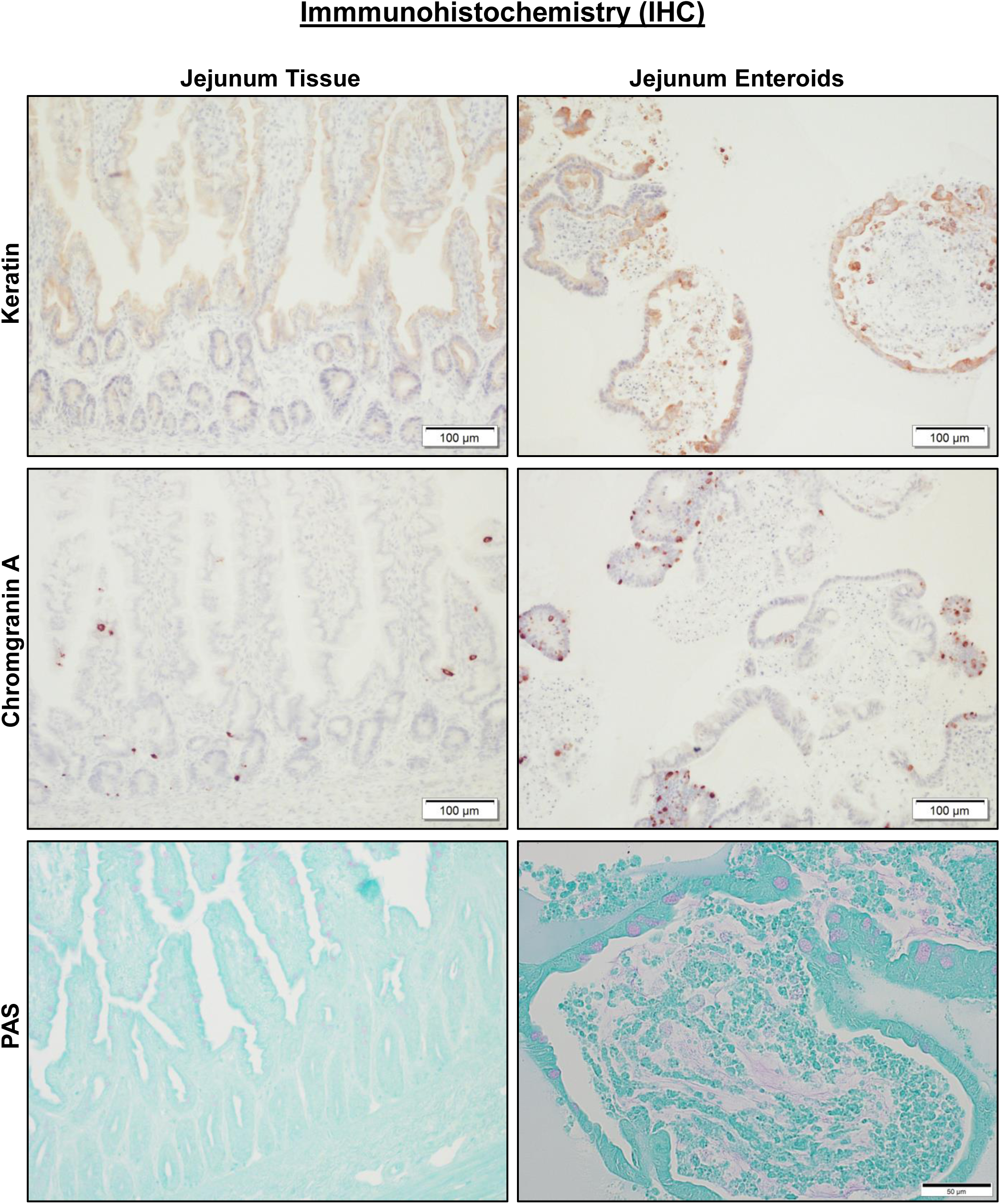
Canine jejunal tissue and enteroids both express markers of epithelial cell lineage. Representative immunohistochemistry (IHC) images comparing staining for marker proteins of epithelial cells and their lineage, including Keratin (epithelial cells, *upper panels*), Chromogranin A (enteroendocrine cells, *middle panels*) and PAS (goblet cells, *lower panels*) on both intact whole jejunal tissue and jejunal enteroids. Tissue and enteroids were counterstained with Hematoxylin (*upper* and *middle panels*) or Alcian Blue (*lower panels*).

### RNA *in situ* hybridization in enteroids

Next, the novel RNA *in situ* hybridization (RNA-ISH) technology (RNAscope) was used to further characterize different intestinal stem cell populations secondary to epithelial cell differentiation, since there are no canine-specific antibodies available for many epithelial cell-line specific markers. RNAscope has a unique probe design strategy that allows for simultaneous signal amplification and background suppression to achieve single-molecule (mRNA) visualization, enabling identification of mRNA expression within cells and in the context of tissue architecture [29]. RNA-ISH shows spatial distribution and cell localization patterns of mRNA expressions, which is critical to identify cell types and their regional location, unlike quantitative PCR, which only provides aggregate mRNA expression. To study intestinal stem cell maturation, we used either early undifferentiated enteroids (3 day old) or differentiated late enteroids (9 day old). ISCs contained within enteroid crypts were identified with canine specific probes for Leucine-rich repeat-containing G-protein coupled receptor 5 (LGR5) and SRY-related HMG-box Sex determining region Y (SRY)-related high-mobility group (HMG) box 9 (Sox9) [30–32]. LGR5, a seminal marker for adult intestinal stem cells [30–31], was observed mainly in the enteroid crypt region of the whole jejunal crypt base (Fig 5a) and quantitated (Fig 5c). As expected, LGR5 expression was significantly reduced (P=0.0001) in the villus portion of both enteroids and full thickness jejunum (Fig 5a, c) as compared to the crypt, confirming that canine intestinal stem cells reside mostly in the crypt area as described in other species [28]. Interestingly, SOX9, a marker for stem cell progenitors [33], was expressed in both crypts and villi of enteroids. However, jejunal crypts had significantly greater SOX9 expression than that observed in their corresponding villi (P=0.0021), indicating that stem cell progenitors reside mainly in crypts. Unlike LGR5, Sox9 expression has also been previously shown at low levels in enteroendocrine and tuft cells, consistent with the low SOX9 expression found in the villi of our dogs [33–34].

**Figure 5.**
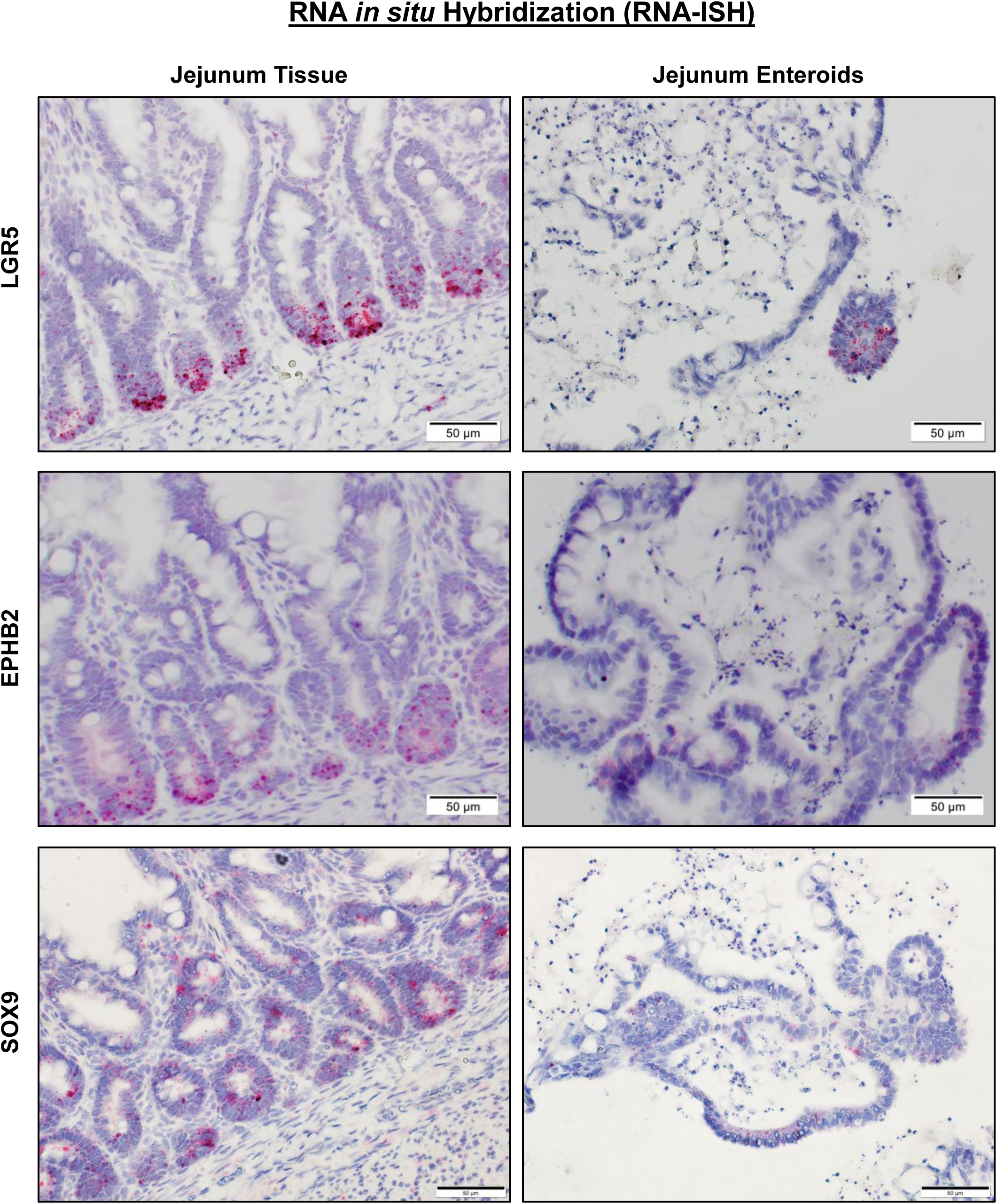
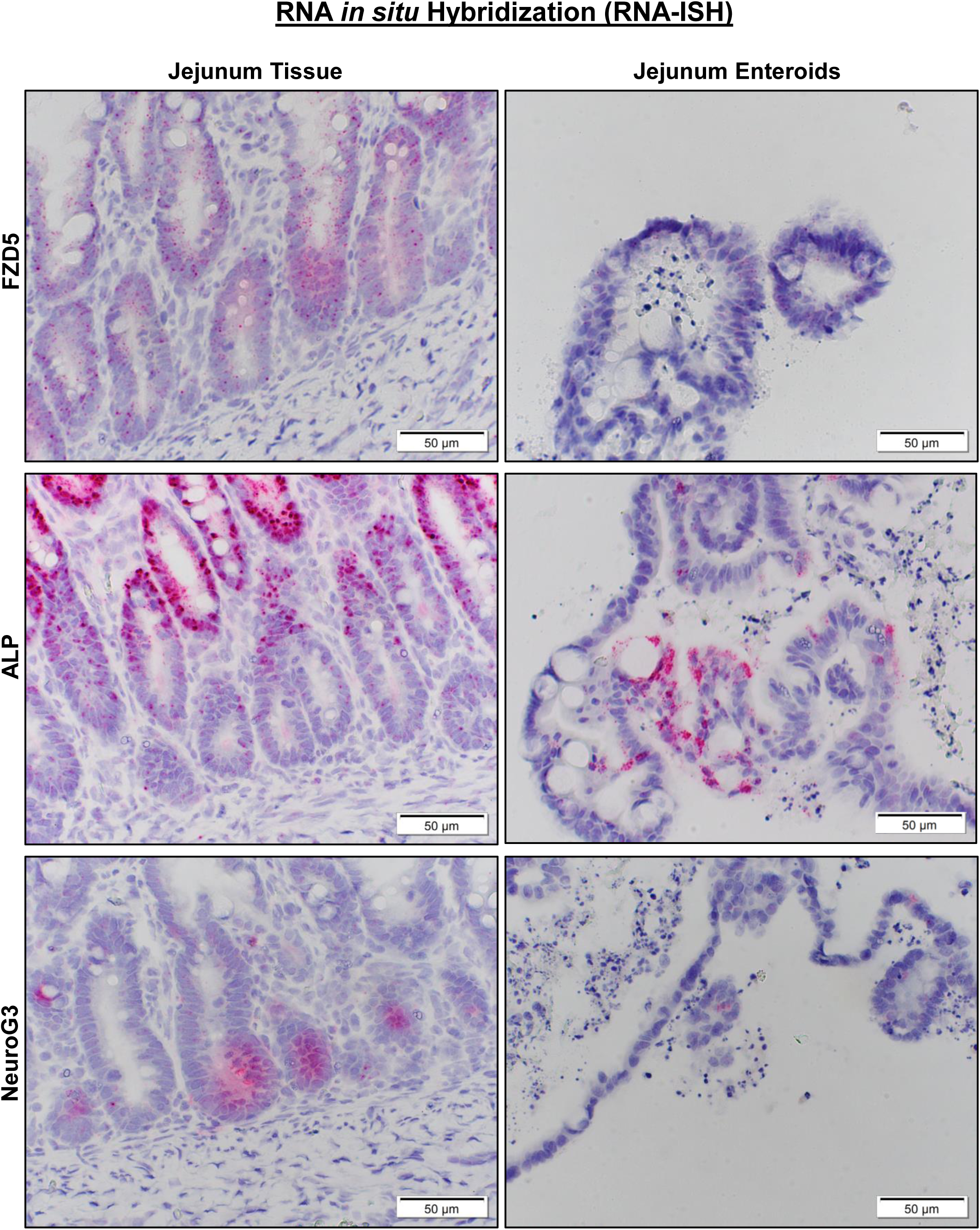
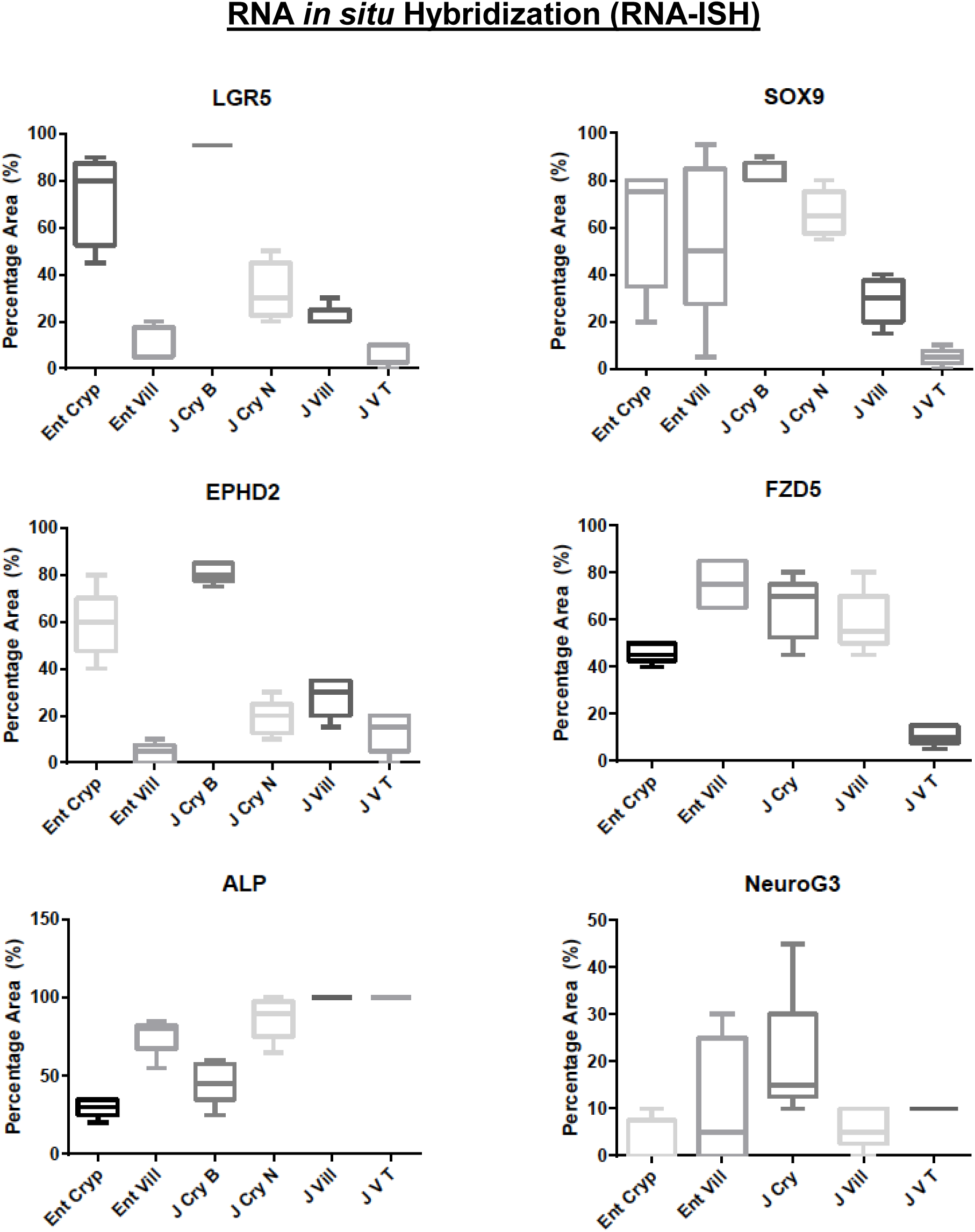
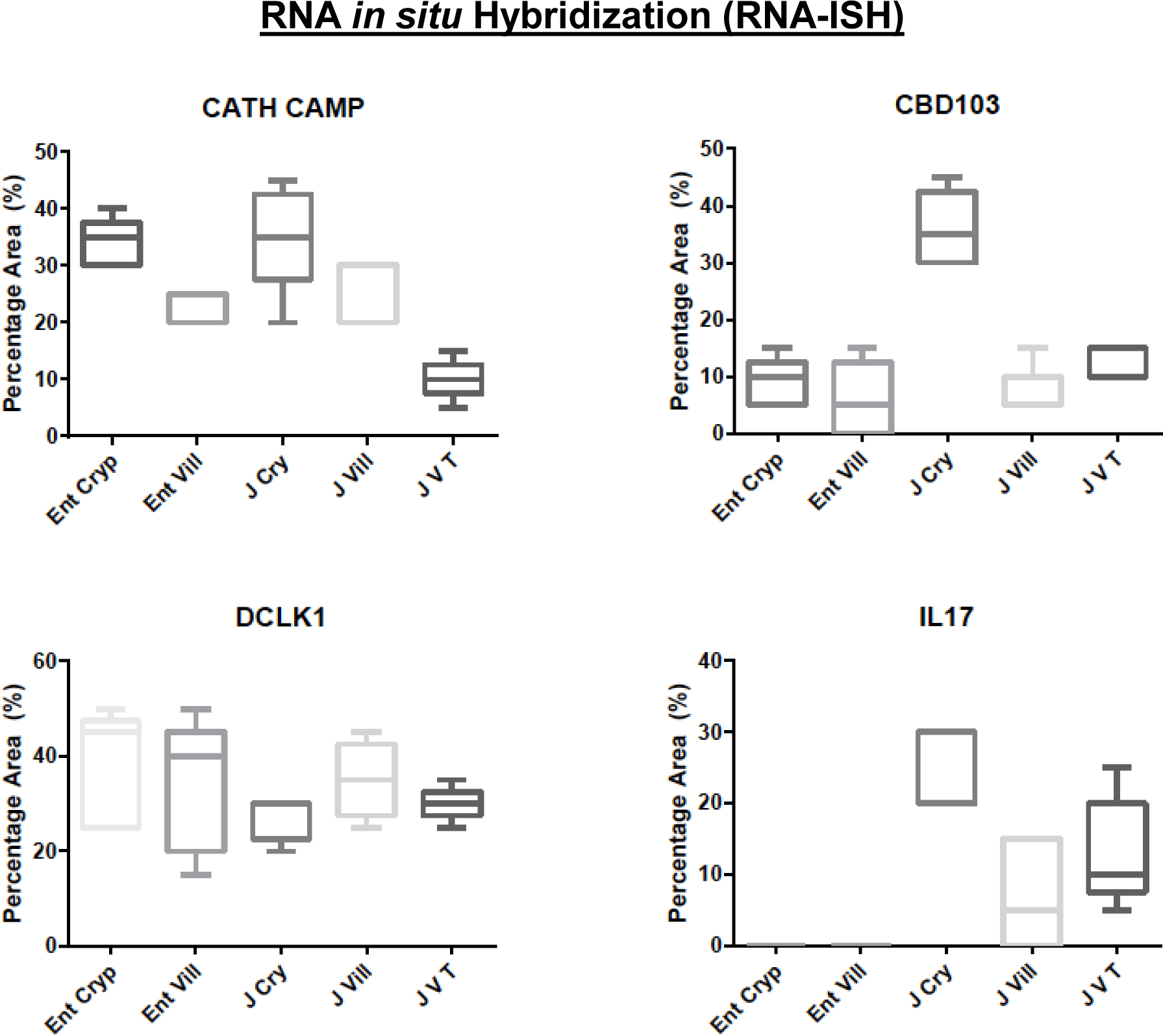
Both canine jejunal tissue and enteroids express mRNA for markers of various epithelial cell lineages. **a** Representative RNA *in situ* hybridization (RNA-ISH) images reveal expression of stem cell markers LGR5, SOX9 and EPHB2 on both intact jejunal tissue and enteroids. SOX9 is also a marker of enteroendocrine and tuft cells, while EPHB2 is also a marker of paneth-like cells. **b** Representative images of gene expression for markers of epithelial lineage, including FZD5 (Paneth like cell), ALP (absorptive epithelium) and Neuro G3 (Enteroendocrine cells) in both intact jejunal tissue and enteroids, as determined by RNA-ISH. **c** Semi-quantitative scoring of RNA-ISH staining (box and whisker plots) for expression of stem cell markers (LGR5, SOX9), paneth-like cell markers (FZD5, EPHB2), absorptive epithelial markers (ALP), and enteroendocrine cells (Neuro G3) in both intact jejunal tissue and enteroids. Specific sites include Enteroid Crypt (Ent Cryp), Enteroid Villus (Ent Vill), whole tissue Jejunum Crypt Base (J Cry B), Jejunum Crypt Neck (J Cry N), Jejunum Villus (J Vill) and Jejunum Villus Tip (J V T). **d** Semi-quantitative expression of paneth cell markers IL-17, CBD 103, and CATH as well as tuft cell marker Dclk1, in both intact jejunal tissue and enteroids, in specific sites as above. Cells and tissue were counterstained with hematoxylin.

Since lysozyme staining for Paneth cells was negative, we next explored expression of other Paneth cell markers responsible for maintaining the ISC niche in dogs [28]. Both ephrin type-B receptor 2 (EPHB2) and Frizzled-5 (FZD5) have a role in canonical Wnt signaling and are also expressed in mouse Paneth cells [35–37]. Similar to LGR5, EPHB2, another marker of intestinal stemness as well as Paneth cells [35], was mainly expressed in the crypts of enteroids and jejunal tissues as compared to the villus compartment (P=0.0001). FZD5 was found both in crypts and, to a lesser degree, in the villi of enteroids and jejunal tissue, with very little expression observed in the villus tips (P=0.001) (Fig 5b-c). Given the putative role of Paneth cells in innate immunity and host defense, we also determined expression of various pro-inflammatory cytokines and antimicrobial peptides produced by Paneth cells, including interleukin-17 (IL-17), beta defensin 103 (CBD 103), and cathelicidin (CATH), using canine specific probes [38]. CBD103 and CATH, both antimicrobial peptides, were observed both in whole tissue and enteroids, whereas IL-17 was found only in whole tissue lamina propria, but not the epithelium (Fig 5d).

We further characterized markers for absorptive enterocyte and enteroendocrine cells with RNAscope target probes for canine Intestinal Alkaline Phosphatase (ALP) and Neurogenin-3 (NeuroG3), respectively [39–40]. Intestinal alkaline phosphatase (ALP), a brush-border enzyme marker for differentiated epithelia, was observed to be more highly expressed in the villi (surface epithelia) of enteroids and jejunal tissue versus cryptal epithelium (P=0.0001) (Fig 5b-c). NeuroG3, a marker for enteroendocrine cells [39] and indirect marker for Paneth cells [40], was dispersed throughout all regions of crypt and villi (Fig 5b-c), with the full thickness tissue jejunal crypt epithelia showing increased expression as compared to the enteroid crypt epithelia (P=0.0296).

Tuft cells, named for their microvilli projections, were identified by a canine Doublecortin-Like Kinase 1 (Dclk1) probe [41]. Tuft cells function as chemosensory cells that initiate type 2 immune responses to helminth parasites [42]. Indeed, under normal conditions the tuft cell population is low, but tuft cell numbers are greatly increased by parasite infection [42]. Dclk1, a marker for tuft cells, was uniformly expressed in crypt and villi of both enteroids and full thickness tissue (Fig 5d).

Finally, we examined the expression of the EP4 prostaglandin-receptor (EP4R), a receptor that has been implicated in the pathogenesis of IBD and is a target of anti-inflammatory and analgesic drugs including nonsteroidal anti-inflammatory drugs (NSAIDS), COX-2 inhibitors, and the piprant class, which are selective EP4R inhibitors. We characterized the expression of EP4R in whole intestinal tissues and enteroids of healthy and diseased dogs to determine the role of EP4R in IBD pathophysiology. There was no difference in epithelial expression of EP4R between biopsies of dogs with IBD and enteroids from IBD dogs (p= 0.37) (Fig 6), indicating that canine enteroids are accurately model EP4R expression in whole tissue. Furthermore, there was no statistically significant difference in EP4R between enteroids of healthy dogs versus enteroids of IBD dogs (p=0.84), nor in the biopsies of healthy dogs vs. dogs diagnosed with IBD (p=0.37) (Fig 6).

**Figure 6.**
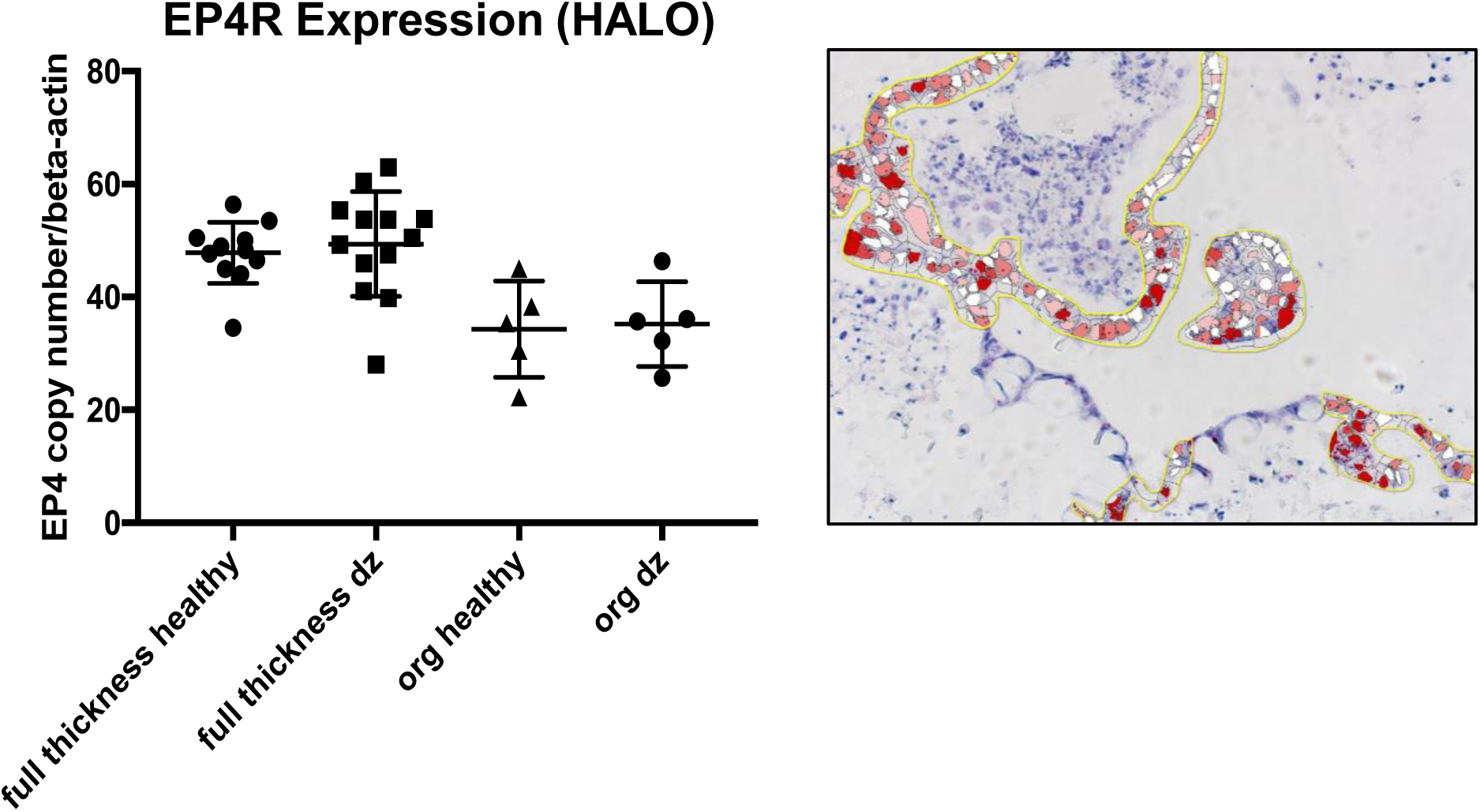
Prostaglandin E2 receptor 4 (EP4R) expression does not change between jejunal tissue and enteroids from healthy and IBD dogs. Representative RNA-ISH image illustrates how the EP4R staining was marked for quantification using Halo software. Box and whisker plot compares the EP4R expression among biopsy tissues (full thickness) and enteroids (org) obtained from both healthy and IBD (dz) dogs.

### Functional assays in canine enteroids

Optical metabolic imaging (OMI), a new technology consisting of fluorescence imaging using a multi-photon microscope, has high resolution and sensitivity to accurately measure cellular metabolic changes, unlike other imaging techniques such as FDG-PET [43]. We used OMI to characterize the metabolic differences between enteroids during differentiation by calculating the optical redox ratio per enteroid. As expected, the optical redox ratio of 7 day old enteroids was approximately twice that of 4 day old enteroids, indicating a lower redox state of the early enteroids versus the older and more differentiated enteroids (Fig 7).

**Figure 7.**
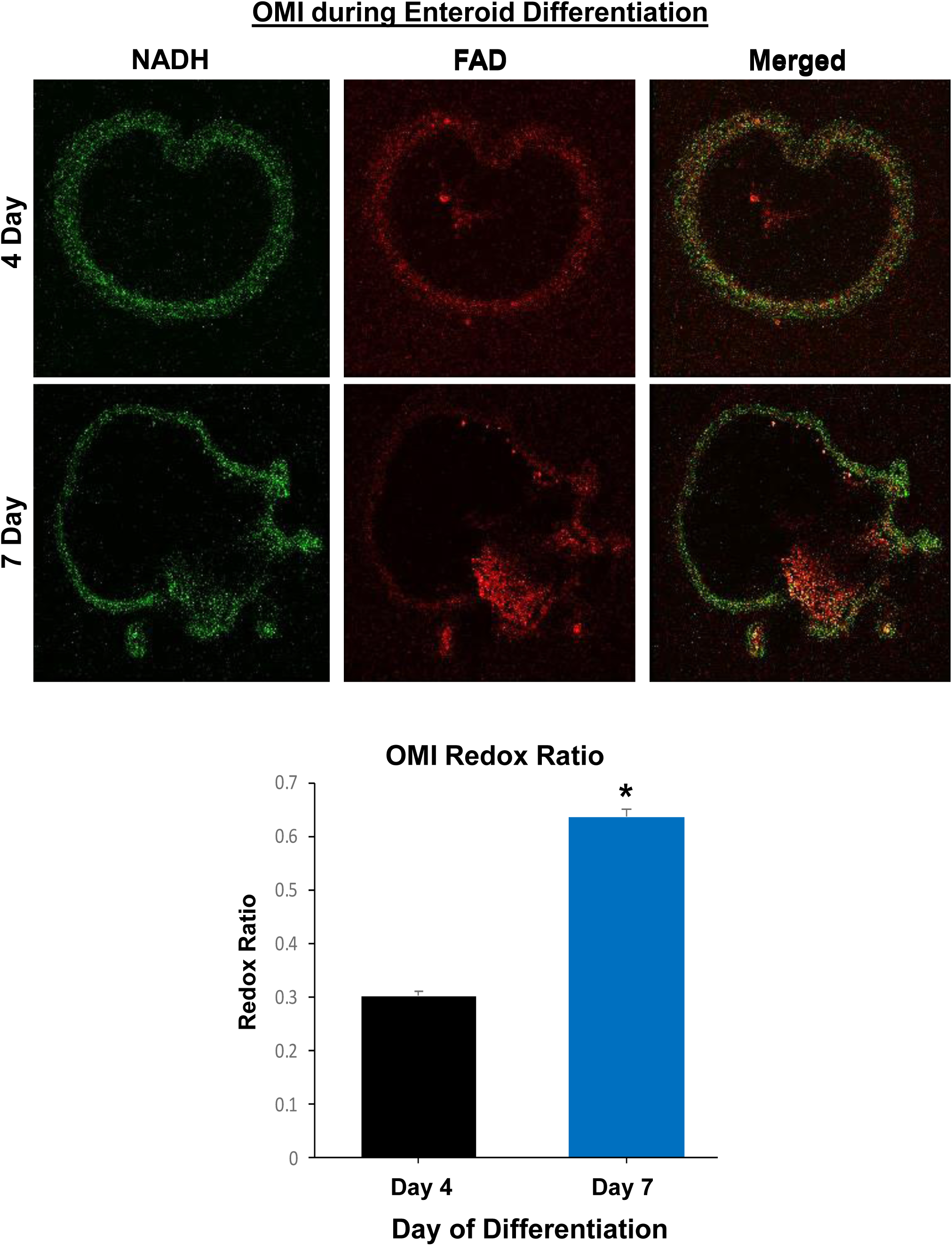
Optical Metabolic Imaging (OMI) reveals metabolic differences between days 4 and 7 of differentiation of canine enteroids. Representative fluorescent images from 4 and 7 day old enteroids are shown. The green image indicates the NADH measurement whereas the red image indicates the FAD, with the merged images on the right. The graph shows the optical redox ratio of 4 day old versus 7 day old enteroids (0.303+0.008 versus 0.637+0.013, respectively, for mean+ SEM).

We next used forskolin, a cyclic adenosine monophosphate (cAMP) agonist, to activate the cystic fibrosis transmembrane conductance regulator (CFTR) chloride channels in intestinal epithelial cells to induce swelling of canine enteroids [13]. As activation of the CFTR chloride channel depends on intracellular cAMP levels, forskolin induces enteroid swelling by increasing water flux into the lumen of the enteroids. This assay can therefore be used as an indirect measure of CFTR function in enteroids [13]. Similar to human intestinal colonoids, incubation with 10 μM forskolin stimulated the swelling of day 2 jejunal enteroids from healthy dogs [13]. Forskolin-induced enteroid swelling was observed after 1, 4, and 24 hours, and the swelling increased in a time-dependent manner (Fig 8), confirming functionality of CFTR in the canine enteroids, similar to intact tissue.

**Figure 8.**
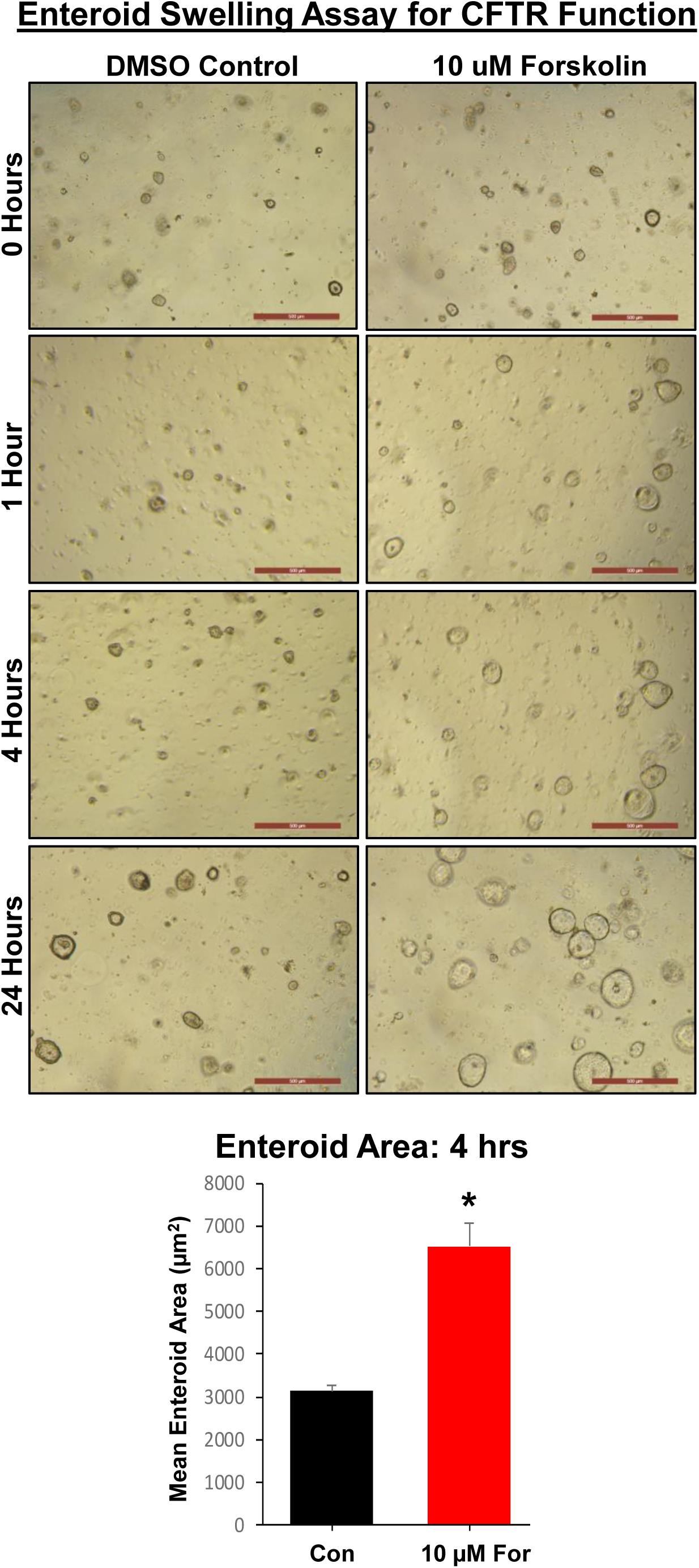
Forskolin induces swelling of canine jejunal enteroids in a time-dependent manner, indicating the presence of functional CFTR. Enteroids were passaged and seeded in matrigel into 24-well plates. After 2 days, enteroids were incubated in media containing vehicle control (DMSO) or 10 μM forskolin. Representative images of enteroids were taken after 0, 4, and 24 hours, at 5X magnification.

We next investigated whether exosome-like vesicles from *Ascaris suum* nematodes could be phagocytized by enteroids. Enteroids incubated for 24 hours with exosome-like vesicles labeled with PKH67 dye demonstrated green fluorescent-labeled exosomes within epithelial cells and within the enteroid lumen (Fig 9). In contrast, enteroids treated with PKH67 dye alone had only DAPI nuclear staining (Fig 9). These data indicated functional uptake of exosomes with transport of vesicles through the epithelial cells and into the enteroid lumen within 24 hours.

**Figure 9.**
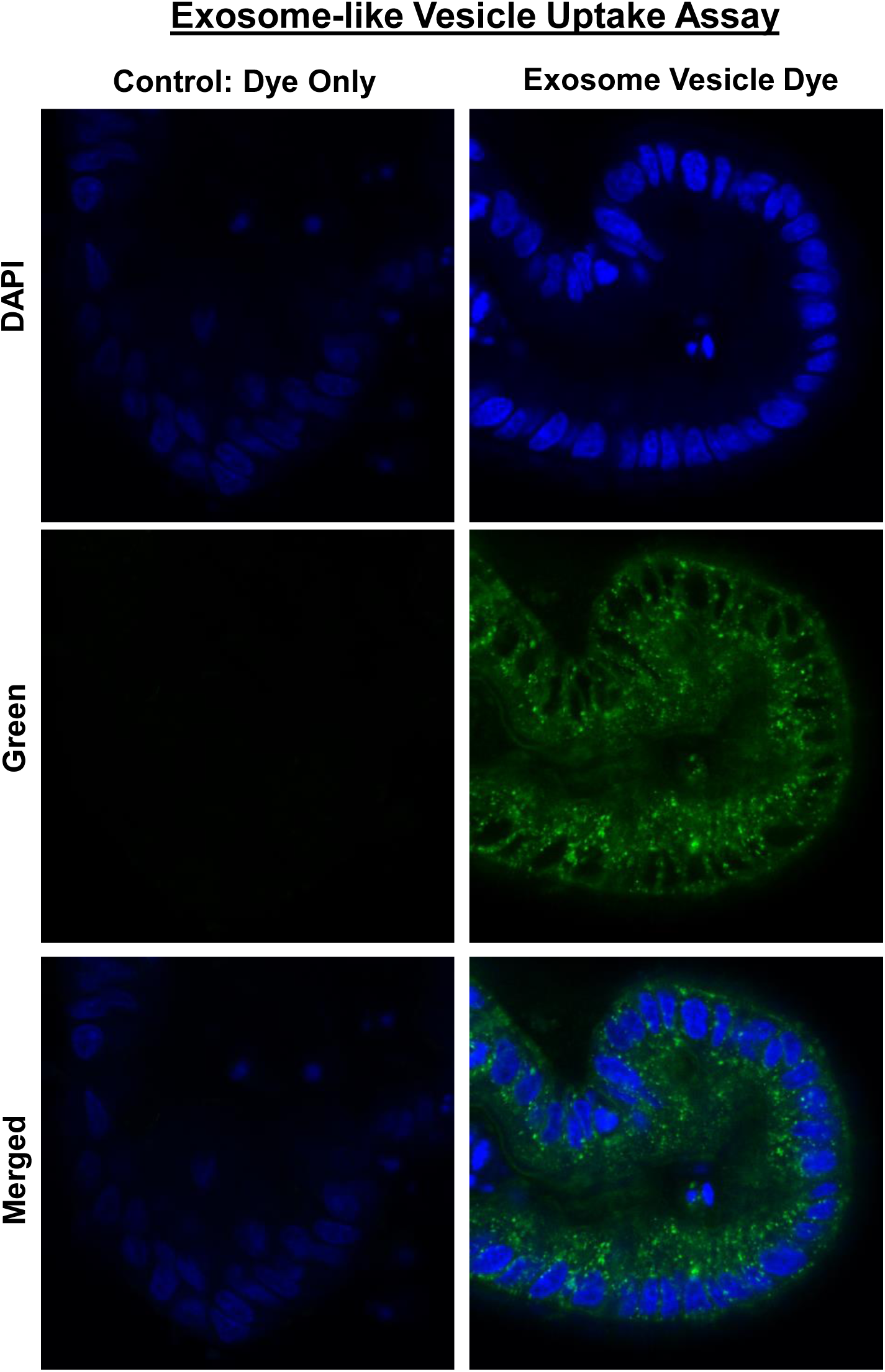
Canine enteroids uptake exosome-like vesicles secreted from the parasite *Ascaris suum.* Representative confocal microscope images taken after 24 hours incubation with control fluorophore alone (green fluorescent PKH67) or exosome-like vesicles (EV) labeled with PKH67 green fluorescent dye. Enteroids were counterstained with the nuclear marker DAPI (blue fluorescence). Only exosome+ PKH67 group demonstrated green fluorescence within epithelial cells and within the organoid lumen. The merged image shows the intracytoplasmic localization of the PKH67 green fluorescent dye labelled EV obtained from *Ascaris suum.*

## Discussion

### Canine enteroids and colonoids as a translational model for GI research

In our study, we were able to successfully culture crypt epithelium obtained from different canine intestinal regions, including the duodenum, jejunum, ileum and colon, using an adapted protocol for isolating human crypt cells [25]. We investigated cultivation of organoids (enteroids and colonoids) from both whole tissue and endoscopically-derived intestinal specimens obtained from healthy dogs and dogs with spontaneous gastrointestinal diseases, including IBD and colorectal cancer. In addition, we could expand intestinal enteroid and colonoid epithelial culture *ex vivo* utilizing a small number of isolated crypts, which allows for the *in vitro* testing of therapeutic drugs and future applications in regenerative medicine. Jejunal enteroids were characterized in depth using IHC, RNA-ISH, morphometric and TEM ultrastructure. Further, we demonstrated the functional utility of canine enteroids by performing OMI, a CFTR function assay and visualization of the active uptake of parasitic extracellular vesicles obtained from *Ascaris suum.* Therefore, these studies form the foundation for the development of canine enteroids as an *ex vivo* large animal model for translational gastrointestinal research.

Canine enteroids and colonoids are a powerful translational and mechanistic tool for identifying molecular targets and therapeutic cures for chronic intestinal diseases such as IBD and CRC. As enteroids and colonoids are composed of epithelial cells alone, they can be used as a platform for the screening of drug candidates that target the epithelial components of intestinal diseases [4]. In addition, canine enteroids and colonoids complement animal-based GI toxicology studies, and potentially reduce the number of animals needed for *in vivo* studies. Canine enteroids and colonoids are also useful for targeted therapy and personalized medicine, as these organoids are derived from individual dogs with different genotypes, environmental risk factors, and drug sensitivity profiles. Combining canine enteroids and colonoids with microfluidic chips will help in the development of precision medicine and the study of interactions between the gut and microbiome. In addition, orthotopic transplantation of canine enteroids or colonoids has the potential to repair or replace damaged or dysfunctional epithelial tissues for regenerative medicine therapies in diseases including IBD, cystic fibrosis, and tuft cell enteropathy.

Among the large animal models used in translational GI research, the dog is particularly relevant as it shares similar environmental, genomic, anatomical, intestinal physiologic and pathologic features with humans [44–47]. Although pigs are by far the most popular large animal model used for transplantation and cardiovascular studies, there are several limitations of the porcine model, including the absence of naturally occurring GI diseases analogous to humans, the high cost for husbandry, the technical difficulties in handling pigs and issues in collecting biological specimens, specifically with regard to obtaining endoscopic biopsies of the GI tract [4].

Undoubtedly, the use of mouse stem cells and organoids revolutionized the study of developmental biology, however, murine models often lack key clinical signs or pathological changes representative of complex human intestinal diseases, such as IBD [48]. In fact, there are growing concerns from the NIH, pharmaceutical industry, and biomedical researchers about the ability of murine models to accurately represent human phenotypes, as illustrated by recurrent failures in translating drug safety and efficacy data from murine models to human patients [49–51]. As such, the success of therapeutic approaches based on intestinal stem cells requires refinement in animal models, including animals that have organs comparable in size and physiology to those of humans [2]. Therefore, it is imperative to develop innovative and robust large animal systems to support intestinal stem cell translational research and the development of new therapeutic discoveries for human digestive diseases [48].

### Canine enteroids model canine intestinal tissue phenotypes

While we obtained enteroids and colonoids from healthy and diseased dogs, we characterized enteroids from healthy jejunum in greater depth due to their importance in nutrient and drug absorption [52]. Our data indicate canine jejunal 3D enteroids are capable of recapitulating the anatomical and physiologic features of native jejunal tissue. Unlike whole intestinal tissues that contain epithelial, mesenchymal and immune cells, adult ISC-derived enteroids and colonoids from mice and humans consist only of epithelial cell populations [30–31]. Epithelial markers (keratin, chromogranin A, and PAS) were present in both canine enteroids and whole jejunal tissue, whereas mesenchymal (vimentin and actin) and immune cell (c-kit and CD3 T cell) markers were found only in whole jejunal tissues (Table 2), indicating that canine enteroids also consist of epithelial populations without other cell types.

Epithelial tight junctions are critical for maintaining the intestinal barrier as well as for a multitude of physiological functions of intestinal epithelial cells, including nutrient absorption and drug transport across the mucosa [53]. In this study, we utilized TEM to assess the structural integrity of epithelial tight junctions as it provides easier characterization of ultrastructural changes compared with other methods, such as IHC [54]. Our TEM studies confirmed that both jejunal tissues and enteroids contain adherens junctions (AJ), tight junctions (TJ) and desmosomes. These proteins are specialized membrane structures that mediate cell-to-cell contact and provide the structural basis for interactions between adjacent intestinal epithelial cells [55–56]. We found that the spacing or gaps in these structures between cells decreased throughout differentiation in the enteroids, creating a less porous intestinal barrier after 6 to 9 days of culture.

The identification of stem and progenitor cell populations in enteroids has proven critical to understanding whether stemness of whole jejunal tissue is retained in enteroids, as well as the impact of *in vitro* culture and maintenance on intestinal stem cells in enteroids. We used a canine specific LGR5 RNAscope probe to identify crypt base columnar stem cells, since LGR5-positive adult ISCs are essential to develop the characteristic crypt-villus enteroid structures in the absence of a non-epithelial cell niche [30]. LGR5 positive adult ISCs were mainly observed in the crypt region of both enteroids and jejunal tissue, which is similar to reports of LGR5-positive adult ISCs in mice and humans [30–31]. In contrast, Sox9, another marker of ISCs, was found to be expressed both in the intestinal crypts and, to a lesser extent, the villus compartments of normal dogs. Although Sox9 is primarily expressed in ISCs and transit-amplifying progenitor cells, it is also found in other epithelial populations, such as enteroendocrine and Tuft cells [32–34]. Therefore, the distribution pattern of SOX9 mRNA in enteroids may not only represent ISC stem cells, but likely indicates the presence of other secretory-lineage epithelial cells. In addition to Sox9, Dclk1 expression in enteroids and intestinal tissues confirms the presence of tuft cells [41].

Positive IHC staining for Alcian Blue/PAS and Chromogranin A for cells in jejunal enteroids indicate that canine ISC can differentiate into specialized epithelial cells including goblet and enteroendocrine cells, as in jejunal tissues. TEM ultrastructure characterization supported the IHC findings and confirmed the presence of enteroendocrine and goblet cells in 3D enteroids. We further investigated the presence of differentiated epithelial cells in enteroids using RNA-ISH, and found expression for the absorptive enterocyte marker ALP and the enteroendocrine/secretory lineage cell marker NeuroG3. In addition, TEM revealed microvilli in the canine enteroids and jejunal tissue, and the number and lengths of microvilli increased during enteroid differentiation. Microvilli are cellular membrane protrusions of absorptive enterocytes containing different populations of brush border enzymes involved in absorption, secretion and cellular adhesion [26]. Our TEM results showing microvilli support our RNA-ISH data of expression of the brush border enzyme ALP, and both microvilli and ALP expression indicate the presence of differentiated absorptive enterocytes.

Paneth cells are highly specialized epithelial cells located in the base of the crypts of Lieberkühn of the small intestine of some species including mice and humans, and are involved in the production of growth factors necessary for ISC function [38]. Since Paneth cells are reported to be absent in some mammalian species, including the dog, functionally equivalent cells remained to be identified in this species. Consistent with this, lysozyme IHC staining and IL-17 RNA-ISH was negative in our canine enteroids as well as the epithelium of full thickness jejunum. We therefore investigated EPHB2 and FZD5 expression in canine enteroids as both receptors are involved in maintenance of intestinal stemness through canonical Wnt signaling and are expressed in murine Paneth cells [35–36, 57]. In addition, the Wnt receptor Fzd5 is required for Paneth cell differentiation and localization to the crypt base in mice [36]. In our study, EPHB2 and FZD5 were expressed both in enteroids and jejunum tissue, suggesting a similarly important role of Wnt signaling in the canine species. In addition to their role in supporting the growth of ISCs, Paneth cells also play an important role in innate immune defense by producing antimicrobial peptides, including CBD 103 and CATH [38]. Our findings of CBD 103 and CATH expression, in addition to EPHB2 and FZD5, support the presence of a functional array of Paneth-like cells in dogs. Altogether, these findings demonstrate that our canine 3D enteroid model system accurately reproduces the anatomical and phenotypical features of canine small intestines, including a broad spectrum of fully differentiated epithelial cells.

Canine enteroids are also a useful *in vitro* model for diseased small intestine tissue. PGE2 is a key mediator of inflammation that acts through EP1, EP2, EP3 and EP4 receptors and has been implicated in the development and progression of various cancers [58–59]. Newer NSAIDs (piprant class) have been developed by targeting EP4R specifically, which should help reduce common GI side effects of NSAID drugs [59]. Our results show that EP4R was expressed in the epithelium of full thickness jejunum and enteroids of healthy and diseased dogs and confirm the utility of canine enteroids to investigate the effects of NSAIDs on EP4R expression in GI tissues. These data also lay the foundation for development of drug testing assays for the piprant class of drugs including grapiprant, a selective EP4R antagonist, using canine enteroids from healthy dogs and dogs with intestinal inflammation, including IBD.

### Functional uses and translational applications of canine intestinal enteroids

In our study, OMI was able to distinguish and quantitate the cellular metabolism between young growing enteroids (Day 4) and differentiated late enteroids (Day 7) using optical redox ratio calculations. OMI has been used previously to determine metabolic activity of various types of tumor organoids in response to drug treatment, but this technique may also prove useful for personalized medicine using canine intestinal and CRC organoids [43]. In addition, we further explored canine enteroid function by examining forskolin-induced swelling of enteroids as a measure of CFTR function. CFTR is the chloride channel mutated in cystic fibrosis, and loss of CFTR function leads to progressive decreases in lung function, pancreatic exocrine dysfunction and intestinal obstruction or constipation [60]. We found that incubation with forskolin doubled the area of canine jejunal enteroids after 4 hours, indicating the presence of functional CFTR chloride channels, similar to human intestinal colonoids [13]. Thus, the forskolin-swelling CFTR function assay with canine enteroids has translational applications in drug screening, especially given that dogs have naturally occurring CFTR mutations similar to humans [13, 61].

Finally, to evaluate the absorptive and barrier functionality of enteroids, we developed a 3D canine enteroid uptake assay using parasitic EVs produced by intestinal helminths. We used EVs from the *Ascaris suum* parasite because of its zoonotic nature and significance in veterinary medicine [62]. EVs are known to elicit host immune responses because of their rich miRNA and bioactive protein contents [63–64], which are hypothesized to induce tolerance towards the helminths in the host organism. Therefore, given the uptake of EVs by our canine enteroids, our 3D enteroid model may be useful to study host-pathogen interactions for parasites that are important in both animal and human disease.

### Canine intestinal crypt isolation, culture and maintenance

Unlike previous reports on canine enteroid culture using collagenase digestion for crypt epithelial isolation [65], we employed a cold EDTA chelation method. This technique is the method of choice for crypt isolation, allowing maximum purity of crypt epithelium and minimum contamination of other cell types [25]. In our study, we used 5-10 times higher EDTA concentration than reported for the mouse crypt epithelial isolation protocol, but similar to that used for humans [25, 66]. In addition, canine ileal tissue samples required a greater EDTA concentration with longer incubation periods for optimal digestion, similar to the treatment of human ileal tissue samples [25]. Variations in EDTA concentrations and the incubation duration for the different intestinal segments are driven by differences in length, numbers and size of villi between various mammalian species [66]. Our data suggest that the canine gut is morphologically more similar to that of humans as compared to the mouse with respect to crypt isolation for enteroid/colonoid culture.

For our studies, we used human ISC culture media containing Wnt-3a for cultivating canine intestinal enteroids and colonoids, as it resulted in better colony forming efficiency (CFE) as compared to mouse ISC culture media, which does not contain Wnt-3a (data not shown). Our observation is consistent with a previous report showing that Wnt-3a, R-spondin-3, and Noggin are required to propagate intestinal enteroids from both farm and companion small animals, including the dog [65]. Additionally, we used Y-27632, an inhibitor of ROCK, and CHIR9902, an inhibitor of GSK-3, for the first 2 days of culture to enhance the survival of canine ISC and to prevent dissociation-induced apoptosis (anoikis). Moreover, the addition of the ROCK inhibitor, Y-27632, into the initial ISC culture medium improved CFE of organoids in our preliminary experiments (data not shown).

## Conclusions

In summary, we have developed and maintained long-term canine enteroid and colonoid 3D models using whole tissue and biopsy samples from healthy dogs and dogs with chronic enteropathies. Our enteroid system complements existing preclinical animal models and provides a translational platform for drug permeability, efficacy, and safety screening. Finally, the tools developed and validated in our lab for the canine 3D intestinal enteroid and colonoid systems include a comprehensive set of reagents and probes, and these will serve as a foundation for using the dog as a translational model for precision and regenerative medicine.

## Methods

### Animals

The collection and analysis of intestinal tissues and biopsies from healthy dogs and dogs with IBD was approved by the Iowa State University (ISU) Institutional Animal Care and Use Committee (IACUC protocols: 4-17-8504-K, 9-17-8605-K, 3-17-8489-K, and 12-04-5791-K).

### Collection of full-thickness and endoscopically-obtained intestinal tissues

For full-thickness intestinal tissues, a 5-10 cm length segment of the proximal jejunum of healthy dogs was collected within 30 minutes of euthanasia. The tissue was next cut open longitudinally and luminal contents removed by forceps. Tissue was immediately placed in wash medium (PBS with 2 mM N-acetylcysteine) and vigorously shaken 10-15 times, repeating washes four times, in order to remove excess mucus, residual luminal contents and other debris [67]. After washing, cleansed tissues were transferred to culture media without growth factor (CMGF-consisting of Advanced DMEM/F12 (Fisher) supplemented with 2 mM GlutaMax-1 (Fisher Scientific), 10 mM Hepes and 100 ug/mL Primocin (InvivoGen)) and incubated on ice. GI endoscopy biopsy forceps (Olympus America) were used to collect mucosa tissue samples from the whole tissue segment. Similarly, 10-15 duodenal, ileal and colonic endoscopic biopsies were obtained by forceps from healthy or IBD dogs under general anesthesia by veterinary gastroenterologists. For clinical research cases, collected biopsies were placed in CMGF-medium on ice and subjected to mechanical cleansing as described above.

### Crypt cell isolation and enrichment from intestinal tissues and biopsies

Epithelial crypts containing adult ISC were isolated and enriched from intestinal tissue samples following a standard procedure adapted from a published human organoid culture protocol [25]. Both whole tissue samples and endoscopic biopsies were cut into small pieces (1-2 mm thickness) with a scalpel and washed 6 times using complete chelating solution (1X CCS) (See Supplementary materials). Pipettes and conical tubes were pre-wetted with 1% Bovine Serum Albumin (BSA) throughout the procedure to prevent adherence of the crypt epithelium to tubes and pipettes, thereby minimizing loss of crypt cells [67]. Then, tissues were incubated with 1X CCS containing EDTA (20mM-30mM) for 45 to 75 minutes at 4°C on a 20 degree, 24 rpm mixer/rocker (Fisher). Length of time for digestion varied depending on collection site, with ileal mucosa requiring 75 minutes of incubation and higher EDTA concentrations (30 mM) versus other intestinal segments. The incubation time was determined by the absence of crypts in tissue fragments, apparent as ‘holes’ and the presence of cellular, dense spheroids under phase contrast microscope (Supplemental Fig. 1) [67]. After EDTA chelation, release of the cryptal epithelium was augmented by trituration and/or mild vortexing in CCS. Additional trituration and/or mild vortexing was carried out after adding fetal bovine serum (FBS, Atlanta Biologicals) to maximize crypt release. After tissue fragments settled to the bottom of the tube, the crypt containing supernatant was transferred to a new conical tube, then centrifuged at 150 g at 4°C for 5 minutes. After centrifugation, the pellet was washed with 10 mL CMGF-medium (see Supplementary materials section) and centrifuged at 70 g at 4°C for 5 minutes. The crypt pellet was then resuspended in 2 mL CMGF-medium and the approximate number of crypts isolated was calculated using a hemocytometer.

### ISC subculture and organoid maintenance

Approximately 400-800 crypts per 10-20 biopsy samples were obtained depending on the GI collection site. Among the four different sites, colonic biopsies yielded the greatest number of ISCs (~800 crypts), the duodenum yielded the least (~400 crypts) and the jejunum & ileum yielded an intermediate number of crypts (~600 crypts). An estimated 50-100 crypts were seeded per well in 30μL of Matrigel (Corning^®^ Matrigel^®^ Growth Factor Reduced (GFR) Basement Membrane Matrix) into a 24 well plate format and incubated at 37°C for 10 minutes [66]. Complete medium with ISC growth factors (CMGF+) and supplemented with 10 μM rho-associated kinase inhibitor (ROCKi) Y-27632 (StemGent) and 2.5 μM glycogen synthase kinase 3β (GSK3β) inhibitor CHIR99021 (StemGent) was added and the plate was incubated at 37°C. The concentration of these growth factors and the requirement of Wnt3a was optimized in our pilot experiments (see Supplementary materials section). The CMGF+ medium with ROCK and GSK3β inhibitors was used for the first 2 days of ISC culture to enhance ISC survival and prevent apoptosis [68]. The enteroid and colonoid cultures were replenished with CMGF + medium every 2 days. After 6-8 days, enteroids and colonoids were completely differentiated, showing a luminal compartment, crypt epithelium and villi-like structures along with exfoliation of denuded epithelia into the lumen (Figure-1). Therefore, the ideal passage expansion of canine enteroids and colonoids is carried out every 5-7 days, just prior to epithelial shedding. For passaging, enteroids and colonoids contained within matrigel were mechanically disrupted using cold CMGF-medium and trituration with a needle syringe (23g ¾”) and pelleted by centrifuging at 4°C x 100 g for 5 minutes. The enteroids and colonoids were resuspended in matrigel and cultured as described above.

### Cryopreservation of enteroids and colonoids

For cryopreservation and bio-archiving, recovery cell freezing media (Invitrogen) was used to facilitate recovery of enteroids and colonoids post-thaw [68]. However, we also tested 90% FBS with 10% DMSO freezing media for some enteroids and colonoids, and did not find a difference between this freezing media and the recovery cell freezing media in quality or quantity of enteroids and colonoids recovered after freezing with either preservation media (Supplemental Fig. 2). We opted to preserve whole canine enteroids and colonoids in liquid nitrogen using the commercial recovery cell freezing media to maintain consistency in cryopreservation. For recovery, enteroids and colonoids were thawed in cold CMGF-medium, centrifuged at 100 g for 5 minutes at 4°C, resuspended in thawed matrigel, and seeded into 24 well plates as above.

### Tissue and enteroid fixation and processing for histopathology and TEM

Representative segments of jejunum tissue and enteroids were incubated in 10% formalin and then stored in 70% ethanol for histopathology (H&E), immunohistochemistry (IHC) and RNA ISH. Fixed tissue and enteroids were paraffin embedded and cut in 4 μm sections onto glass slides. For TEM, enteroids and tissue were initially placed in a fixative solution composed of 2% paraformaldehyde/3% glutaraldehyde/0.1 M cacodylate with pH 7.2, followed by 1% osmium tetroxide post fixative. Thick (μm) and ultrathin (50-100 nm) sections were cut by microtome, collected on copper grids and observed under a JEOL 2100 200kV STEM.

### IHC characterization of enteroids and tissues

IHC was performed using established protocols [69–72]. Briefly, sections were deparaffinized and rehydrated using an automated system with manual antigen retrieval and blocking steps. Primary antibody, biotinylated secondary antibody, horseradish peroxidase-streptavidin and NovaRed staining were applied on the section in a sequential order followed by counterstaining. Markers used in IHC were pan-Keratin, Chromogranin A, Vimentin, Actin, c-Kit, T cell, PAS and Lysozyme. Presence and absence of these markers was evaluated using light microscopy and the cell types determined by morphology, location and staining, and pictures were taken. The list of markers and the antibody source, dilution and incubation details are given in Supplementary Table 1.

### RNA-ISH characterization of jejunal enteroids and tissues

RNA *In Situ* Hybridization (ISH) was performed using the RNAscope 2.5 High Definition (Red) kit per the manufacturer’s protocol (Advanced Cell Diagnostics, Newark, CA). Briefly, enteroids and tissue sections were deparaffinized, boiled in target retrieval solution, and incubated in protease buffer. Then sections were hybridized with specific oligonucleotide probes for cell-surface markers of intestinal stem cells, epithelial differentiation and maturation, and then the respective mRNA was serially amplified [29]. The details of the probe and sequences are given in Table 2. After amplification, signal was detected with RED-B and RED-A, and hematoxylin used as counterstain. The expression of mRNA markers was calculated using the singleplex semi-quantitative scoring criteria adapted from ACD Pharma Assay Services data analysis for the intensity of staining (<1 dot/10 cells is no staining; 1-3 dots/cell is +; 4-9 dots/cel is ++; 10-15 dots/cell is +++; >15 dots/cell is ++++). The staining intensity between different mucosal regions including crypt and villi was compared.

### Optical metabolic imaging (OMI) of canine enteroids

Day 4 and 7 enteroids from healthy dogs underwent OMI as one measure of functional activity, using a two-photon FLIM microscope incorporating time correlated single photon counting [43]. NADH and FAD were imaged at two-photon excitation wavelengths of 750 nm and 890 nm, respectively, with emission collected between 400-480 nm and 500-600 nm, respectively, using a 20X/1.15 NA water immersion objective. A minimum of 15 image volumes were captured within each sample. To quantify the effect of different age enteroids (4 day versus 7 day old) on cellular metabolism of the enteroids as a whole, we estimated the optical redox ratio of the intensities of cellular cofactors NADH and FAD, as described by Walsh et al. [43]. Optical redox ratio was calculated as NADH intensity divided by FAD intensity per organoid, and was the average of 15 samples per condition.

### CFTR functional assay using canine enteroids

Enteroids were passaged and seeded in matrigel into 24-well plates as above. After 2 days, enteroids were incubated in CMGF+ media containing vehicle control (DMSO) or the indicated treatments [13]. Representative images of enteroids were taken after 0, 1, 4, and 24 hours, at 5X magnification on an inverted microscope using the Leica Application Suite (LAS) software. 6 wells and 2 fields per well were used for each condition, and the average enteroid area for each field was calculated using ImageJ software.

### Exosome-like vesicle uptake assay using canine enteroids

*Ascaris suum* female adult nematode worms were obtained from JBS Swift and Co. pork processing plant (Marshalltown, Iowa, USA) and cultured using the *Ascaris* Ringers solution (ARS): 13.14 mM NaCl, 9.47 mMCaCl2, 7.83 mM MgCl2, 12.09 mM Tris, 99.96 mM Sodium acetate, 19.64 mM KCl, pH 7.8 supplied with 5 mM Glucose, 10,000 units pen/strep, 10 μg/mL ciprofloxacin and 0.25μg/mL amphotericin [63]. Culture media was collected between 24 h to 48 h and passed through a 0.22 μm filter (EMD Millipore, USA). Extracellular vesicles (EVs) secreted by the helminths were purified by ultracentrifugation protocol as previously described [63]. Briefly, the medium containing EVs was centrifuged at 4°C x 120,000 g for 90 mins (Becman SW32Ti rotor), then the EV pellet was resuspended in DPBS (Thermofisher, USA), transferred to a new tube and centrifuged at 4°C x 154,000 g for 2 hours (Beckman TLA55 rotor). The EVs pellet was resuspended with DPBS and stored at −80°C. The protein concentration of EVs was determined using a Qubit Fluorometer (Thermofisher, USA). Purified EVs were stained with PKH67 Fluorescent dye (Sigma, USA) as per the manufacturer’s protocol. In brief, 10μg EVs were resuspended with the diluents solution, and then stained with PKH67 dye at room temperature for 5 mins. The labelling reaction was stopped by mixing with an equal volume of FBS. Then the EVs were pelleted by centrifuging at 4°C x 154,000 g for 2 hours (Beckman TLA55 rotor). Pellets were resuspended with 1000μL DPBS (Thermofisher, USA). An equal volume of CMGF+ medium was mixed with EVs labelled with PKH67 dye. As a positive control, an equal volume of CMGF+ medium was mixed with PKH67 dye alone. Ileal enteroids from healthy dogs were cultivated on 8 well chamber slides, and on Day 3, enteroids were incubated with CMGF+ medium containing control, dye only control, or labelled exosome-like vesicles for 24 hours at 37°C. Then, enteroids were washed three times with PBS and fixed in 4% paraformaldehyde (Sigma-Aldrich). After fixation, enteroids were washed with PBS, counterstained with DAPI and visualized at 40X magnification using a Leica TCS SP5 X Confocal/multiphoton microscope system (Leica Microsystems Inc., Buffalo Grove, IL).

### Statistical analysis

Data were analyzed using graph pad prism version 7 and R version 3.5. RNA-ISH semi-quantitative data were analyzed using Kruskal-Wallis Statistical test for multiple comparisons of groups. Dunn’s post hoc testing was used wherever specific comparisons between two groups was needed. Student’s t-test was used to analyze significance for other experiments where appropriate. Statistical significance was set at p< 0.05.

## Declarations

### Ethics approval and consent to participate

All animal studies were reviewed and approved by Iowa State University IACUC, as detailed in the Methods.

### Consent for publication

All authors have given consent for publication.

### Availability of data and materials

All data generated and analyzed in this study are included within the article or supplementary materials. Canine intestinal organoids are available upon request.

## Competing interests

JPM, KA, and AJ would like to disclose a competing financial interest and management role in 3D Health Solutions, Inc., an entity that provides 3D organoid testing services. The other authors have no competing interests to declare.

## Funding

This work was supported by a Departmental Research Start-Up Grant at ISU to KA, and by a Miller Research Award from the Office of the Vice-President for Research at ISU to JM.

## Author Contributions

LC and DCB wrote the manuscript. LC, DDK, KA, AJ, and JM were responsible for the concept and design of the overall study and interpretation of data. LC, DDK, TA, and DCB were involved in experimental design, acquisition of data, analysis and interpretation of data. ABM, YMA, YQ, QW, MM, DM, MS, MFZ, and MW provided technical assistance and guidance. WY, MK, NME, and ES provided unique samples and reagents. KA, AJ and JM supervised the project and provided critical revision of manuscript. All authors read and approved the final manuscript.

## Acknowledgments

The authors would like to thank Tracy Stewart and the Roy J. Carver High Resolution Microscopy Facility (HRMF) at Iowa State University, for assistance with confocal microscopy and TEM.

## Author’s information

Not applicable

